# The brassinosteroid biosynthesis gene *TaD11-2A* controls grain size and its elite haplotype improves wheat grain yields

**DOI:** 10.1101/2022.02.17.480859

**Authors:** Huiyuan Xu, Han Sun, Jiajin Dong, Chengxue Ma, Jingxue Li, Zhuochun Li, Yihuan Wang, Junqi Ji, Xinrong Hu, Meihui Wu, Chunhua Zhao, Ran Qin, Jiajie Wu, Fei Ni, Fa Cui, Yongzhen Wu

**Affiliations:** College of Agriculture, Key Laboratory of Molecular Module-Based Breeding of High Yield and Abiotic Resistant Plants in Universities of Shandong, Ludong University, Yantai, Shandong, China; State Key Laboratory of Crop Biology, College of Agronomy, Shandong Agricultural University, Taian, Shandong, China

**Keywords:** *Triticum aestivum*, favorable haplotype, grain size, 1000-grain weight, grain yield per plant

## Abstract

Brassinosteroids (BRs) control many important agronomic traits, therefore the manipulation of BR components could improve crop productivity and performance. However, the potential effects of BR-related genes on yield-related traits and stress tolerance in wheat (*Triticum aestivum* L.) remain poorly understood. Here, we identified *TaD11* genes in wheat (rice *D11* orthologs) that encoded enzymes involved in BR biosynthesis. *TaD11* genes were highly expressed in roots (Zadoks scale: Z11) and grains (Z75), while expression was significantly suppressed by exogenous BR (24-epiBL). Ectopic expression of *TaD11-2A* rescued the abnormal panicle structure and plant height (PH) of the *cpb1* mutant, and also increased endogenous BR levels, resulting in improved grain yields and grain quality in rice. Natural variations in *TaD11-2A* were associated with significant differences in yield-related traits, including PH, grain width (GW), 1000-grain weight (TGW), and grain yield per plant (GYPP), and its favorable haplotype, *TaD11-2A-HapI* was subjected to positive selection during wheat breeding. Additionally, *TaD11-2A* influenced root length and salt tolerance in rice and wheat at seedling stages. These results indicated the important role of BR *TaD11* biosynthetic genes in controlling grain size and root length, and also highlighted their potential in the molecular biological analysis of wheat.

**Highlight:** The brassinosteroid biosynthesis gene *TaD11-2A* regulates grain size and root length and its haplotype favorably improves grain yields and salt tolerance in wheat.

## Introduction

Wheat (*Triticum aestivum* L.) is one of the most important crops in the world. It is highly adaptable and widely distributed, and provides approximately 20% of humans’ calorie and protein quotas. By 2034, it is estimated that wheat production will have to increase by another 50% from current levels to feed increasing global populations (Chen *et al*., 2020). Therefore, current wheat breeding efforts have focused on factors improving grain yield and grain quality, one of which is grain size. To date, several genes regulating wheat grain size have been identified, characterized, and shown to affect grain weight or grain quality; *TaCYP78A3* (Ma *et al*., 2015), *TaGS3* (Yang *et al*., 2019; Zhang *et al*., 2020), *TaGS5* (Ma *et al*., 2016; Wang *et al*., 2015), *TaGW2* (Qin *et al*., 2014; Simmonds *et al*., 2016; Su *et al*., 2011; Yang *et al*., 2012), *TaGW8* (Ma *et al*., 2019; Yan *et al*., 2019), *TaDA1* (Liu *et al*., 2020a), *TaSG-D1* (Cheng *et al*., 2020), and *TaNAC019* (Gao *et al*., 2021; Liu *et al*., 2020b). These wheat gene counterparts are believed to control grain size via different pathways, including G-protein signaling, protein degradation, phytohormone signaling and starch synthesis, or unknown pathways (Gao *et al*., 2021; Ma *et al*., 2015; Zuo and Li, 2014). Nevertheless, few studies have reported grain size regulation via brassinosteroid (BR) biosynthesis in wheat.

BRs are a class of plant steroid hormones that play essential roles in plant growth and development by eliciting a wide range of morphological and physiological responses, and also tolerance to abiotic and biotic stresses (Nolan *et al*., 2020; Wei and Li, 2016). BR biosynthetic pathways have been largely elucidated, and include three pathways leading to the production of C_27_-, C_28_-, and C_29_-types of BR (Bajguz *et al*., 2020). C_28_-BR types are the most widely distributed pathways and include the brassinolide (BL) which display the greatest biological activity (Clouse and Sasse, 1998). Thus far, most BR-related genes, such as *CYP90C1, CYP85A2*, and *DWF4* in *Arabidopsis* (Choe *et al*., 1998; Kim *et al*., 2005; Nomura *et al*., 2005), *BRD1* (also known as *OsBR6ox*), and *OsDWARF4* and *DWARF11* (*D11*) in rice (Hong *et al*., 2002; Mori *et al*., 2002; Sakamoto *et al*., 2006; Tanabe *et al*., 2005) have been characterized via the C_28_-BR biosynthesis pathway. These genes belong to the cytochrome P450 family and affect a wide range of agronomic traits. For example, *DWF4* encodes CYP90B1 which catalyzes the C-22 hydroxylation of campesterol and influences plant architecture in *Arabidopsis* (Choe *et al*., 1998). *D11* (also known as *CPB1*) encodes CYP724B1 which catalyzes the conversion of 6-deoxocathasterone and 3-dehydroteasterone to 6-deoxotyphasterol and typhasterol, respectively, and affects plant height (PH), panicle architecture, and grain shape in rice (Tanabe *et al*., 2005; Wu *et al*., 2016).

Previous studies reported that the genetic manipulation of genes in BR biosynthetic pathways provide considerable potential for improved crop productivity and performance (Nolan *et al*., 2020). In rice, transgenic plants using an *S-adenosylmethionine synthase* to drive *DWF4* expression (derived from rice, maize, or *Arabidopsis*) increased yields by approximately 15%−44% over wild type plants (Wu *et al*., 2008). Targeted expression of *CPB1*, using panicle-specific promoters, enhanced grain size and grain yield per plant (GYPP) (Wu *et al*., 2016). *ZmD11* overexpression significantly increased seed size, seed weight, and seed quality-related traits in rice and maize (Sun *et al*., 2021). The ectopic overexpression of *AtDWF4* in the oilseed crop, *Brassica napus* significantly increased seed yields, improved tolerance to dehydration and heat stress, and resistance to two necrotrophic fungal pathogens (Sahni *et al*., 2016). However, the potential impact of BR biosynthetic genes on wheat yields and stress resistance remains unknown.

In this study, we isolated and characterized three *TaD11* homologs from wheat. Expression analyses showed that the *TaD11* genes were dominantly expressed in roots (Z11) and grains (Z75) and were significantly repressed by exogenous BR (24-epiBL) or the BR synthesis inhibitor, brassinozole (BRZ). Transgenic assays demonstrated that *TaD11-2A* overexpression restored the defective phenotypes of the *cpb1* mutant containing an allelic mutation of *OsD11*, and enhanced grain yield and grain quality by increasing endogenous BR levels. The *tad11-2a* mutant exhibited dwarfism, smaller grains, and 24-epiBL sensitivity. Haplotype analysis revealed that the favorable haplotype, *TaD11-2A* was significantly associated with higher grain width (GW), 1000-grain weight (TGW), and GYPP. Moreover, *TaD11-2A* affected root length and salt tolerance in rice and wheat at seedling stages. These findings suggested that *TaD11’s* are valuable reagents for dissecting grain yield patterns and grain quality balance, and also investigating BR functions in wheat grain and root development.

## Materials and methods

### Plant materials and growth conditions

The rice *cpb1* mutant (Wu *et al*., 2016) and *japonica* cultivar Zhonghua 17 (ZH17) were both used for genetic transformation and functional validation studies. The wheat *tad11-2a* mutant (Kronos2702) was screened from the *Triticum turgidum* cv. Kronos TILLING population, developed by Dr. Jorge Dubcovsky at UC Davis, CA, USA, and used for gene function analysis. These reagents were grown in experimental fields at Ludong University in Yantai, Shandong province, China. The natural population used for haplotype analysis contained 314 wheat accessions, including 250 in Asia, three in Africa, nine in Europe, 42 in North America, four in South America, and six in Oceania, (Supplementary Table S4). These natural group materials were planted in seven different environments (E1–E7) in four regions during the years 2017–2020, using a randomized complete block design with two replications (Supplementary Table S5).

### Phylogenetic tree and conserved domain analysis

Full-length amino acid sequences of *TaD11* and homologs in cereals were retrieved from the Unité de Recherche Génomique Info (URGI) (http://urgi.versailles.inra.fr/) and Gramene (http://www.gramene.org/) databases. Multiple protein sequences were aligned using the ClustalW program in DNASTAR v11.1 (https://www.dnastar.com/). Based on amino acid sequence alignment results, a neighbor-joining tree was constructed in MEGA v10.1 using the Jones-Taylor-Thornton (JTT) model with 1000 bootstraps (Kumar *et al*., 2018). Conserved domains in protein sequences were checked using the National Center for Biotechnology Information (NCBI) Batch Web CD-Search Tool (https://www.ncbi.nlm.nih.gov/Structure/bwrpsb/bwrpsb.cgi).

### RNA extraction and quantitative real-time PCR (qRT-PCR)

Total RNAs were extracted from various tissues using the MiniBEST Plant RNA Extraction Kit (Takara Bio, Beijing, China). First-strand cDNAs were synthesized using TransScript One-Step gDNA Removal and cDNA Synthesis SuperMix (TransGen Biotech, Beijing, China). RT-PCR was conducted using TB Green Premix Ex Taq II (Takara Bio) in a CFX96 Real Time PCR Detection System (Bio-Rad, Hercules, CA, USA). Each set of experiments had at least three biological replicates with at least three technical triplicates. Rice *ubiquitin* and wheat *GAPDH* were used as internal references for rice and wheat, respectively, and gene levels were normalized using the relative quantitative method (Livak and Schmittgen, 2001). Primers were designed by Primer Premier 6.0 (Supplementary Table S1).

### Vector construction, rice transformation, and trait measurements

To generate the overexpression vector, p*Ubi*::*TaD11-2A*, the full-length open reading frame of *TaD11-2A* was generated using PCR and inserted into the modified binary vector, pCAMBIA1301 (Wu *et al*., 2016) using a homologous recombination method. The construct was then transformed into the *cpb1* mutant and ZH17 plants, respectively, using an *Agrobacterium*-mediated approach. Phenotypic measurements of transgenic plants were performed using three independent transgenic lines (15 positive plants/line). GL, GW, and grain area were evaluated using the intelligent test and analysis system (TOP Cloud-Agri Technology, Zhejiang, China). For grain quality-related traits, total starch and amylose levels were determined using the K-TSTA and K-AMYL kits (Megazyme, County Wicklow, Ireland), respectively, according to manufacturer’s instructions, while protein levels were measured according to a previous method (Bradford, 1976).

### Phenotypic evaluation in wheat

Each wheat accession from the natural population was space-planted in a 1.5 m single-row plot, with 5 cm between plants, and 30 cm between rows. Five plants of each accession in each replicate were selected to investigate 15 agronomic traits, including PH, SL, SSL, AL, FLL, SN, SNPS, GNPS, GL, GW, TGW, GYPP, PC, TW, and WGC. Three quality-related traits, including PC, TW, and WGC, were measured using the Antaris II FT-NIR analyzer (Thermo Fisher Scientific, USA). The BLUE values for each trait were calculated using the R package lme4 (Douglas Bates *et al*., 2015). Root phenotypes at seedling stages, including total root length and total root volume, were scanned using the Microtek ScanMaker (Zhongjing Technology, Shanghai, China) and analyzed using the LA-S plant root phenotype analysis system (Wseen Testing Technology, Hangzhou, China).

### 24-Epibrassinolide, chemical treatments, and the quantification of endogenous BRs

For all treatments, 24-epibrassinolide (24-epiBL, Sigma-Aldrich, Shanghai, China) and BRZ (Sigma-Aldrich) were dissolved in absolute ethanol to a stock concentration of 5 mM, and diluted in sterile water to a final concentration of 1 _μ_M. Subsequently, 7-day-old seedlings of the wheat cultivar, Chinese Spring were transferred to a hydroponic tank containing 1 _μ_M 24-epiBL or 1 _μ_M BRZ or mock control (distilled water). The roots from six seedlings were stored in liquid nitrogen at 0 h, 1 h, and 3 h after treatments for *TaD11* expression analysis. For endogenous BR quantification, approximately 2 g shoot tissues at the seedling stage were collected from control and transgenic plants, under hydroponic conditions. Three BRs, brassinolide, castasterone, and 6-deoxocastasterone were measured at NJRuiYuan Co., Ltd. as described previously (Xin *et al*., 2013). Three biological replicates were assayed for each sample.

### Lamina inclination assay

Normally germinated 2-day-old wheat seeds were transferred to hydroponic boxes containing 96 wells, and grown for 5 days in a growth chamber at 23°C, 50% relative humidity, and a photoperiod of 16 h/8 h light/darkness. After this, 2 _μ_l ethanol containing 0, 10, 100, or 1000 ng 24-epiBL (Sigma-Aldrich) was spotted at the top of the lamina of seedlings (Fujioka *et al*., 1998). After 3 days incubation, seedlings were photographed and the angle between the lamina and leaf sheath measured using ImageJ (https://imagej.nih.gov/ij/index.html). Fifteen plants were used for each measurement.

### Salt and drought tolerance assay

For this assay, the culture conditions of wheat seedlings were the same as the lamina inclination assay. Seedlings at the one leaf stage were treated in 1/2 Hogland solution (pH 6.0) containing 1.5% NaCl (Sigma-Aldrich) or 20% polyethylene glycol (PEG) 6000 (Sigma-Aldrich) for 3 or 5 days, respectively. Then, growth conditions were restored to control conditions and plants recovered for 5 days. Seedlings with new leaf growth were counted as surviving plants. The water and nutrient solution was replaced every 3 days. In rice, the evaluation of tolerance to salt and drought at seeding stages was conducted as described previously (Sun *et al*., 2021).

### Population structure and GWAS

Genotyping of the 314 accessions was performed using the Wheat 55K SNP array at Beijing Compass Biotechnology Co., Ltd. In total, 24838 unique SNPs were obtained by filtering with probe sequences BLAST the reference genome (IWGSC RefSeq v2.1) and PLINK1.9 ‘--maf 0.01 --geno 0.2 --mind 0.2’. The population structure was assessed using a subset of 3871 SNPs (LD < 0.4) using ADMIXTURE v1.3.0 (Alexander *et al*., 2009). The R package pophelper was used to generate ancestry barplots (Francis, 2017). We performed a GWAS of 24840 markers (*TaD11* haplotypes and 24838 unique SNPs) and 15 agronomic traits using the GLM approach in TASSEL v5.2.73 (Bradbury *et al*., 2007), with principal components of principal component analysis (PCA) as covariates. The LDBlockShow v1.40 (Dong *et al*., 2021) package was used to draw a LD heatmap.

### Statistical analysis

Two-tailed Student’s *t* tests and Duncan’s multiple-range tests were performed using IBM SPSS Statistics v26 (IBM Corp., Armonk, NY, USA). Significance was accepted at \**P* < 0.05 and \*\**P* < 0.01.

## Results

### Identification and characterization of TaD11 genes

To investigate the functions of *D11* in wheat, we performed BLASTP analyses using the rice D11 (OsD11) protein sequence as a query to search the International Wheat Genome Sequencing Consortium (IWGSC) RefSeq v2.1 database (Zhu *et al*., 2021). We identified three highly homologous genes located on chromosomes 2A (TraesCS2A03G0818100), 2B (TraesCS2B03G0904700), and 2D (TraesCS2D03G0759600), which we designated *TaD11-2A, TaD11-2B*, and *TaD11-2D*, respectively (Fig. 1A). Primers were designed according to assembled sequences (Supplementary Table S1) to clone genomic sequences and cDNAs from the wheat cultivar, Chinese Spring. The genomic length of *TaD11-2A, TaD11-2B*, and *TaD11-2D* sequences were 4050 bp (base pairs), 4144 bp, and 4087 bp, respectively. Each gene consisted of nine exons and eight introns, and encoded 476, 475, and 476 amino acid proteins, respectively (Fig. 1B; Supplementary Table S2). Sequence analyses showed that predicted TaD11 proteins all contained a CYP90-like domain (cd11043) and belonged to the cytochrome P450 superfamily (cl41757) (Supplementary Fig. S1), suggesting similar functions in wheat. Phylogenetic analyses revealed that D11 homologs were divided into two groups in cereals, C3 and C4 clades, and that TaD11 proteins were most closely related to HvD11 (HORVU2Hr1G081650) from *Hordeum vulgare* (Fig. 1A).

**Fig. 1.**
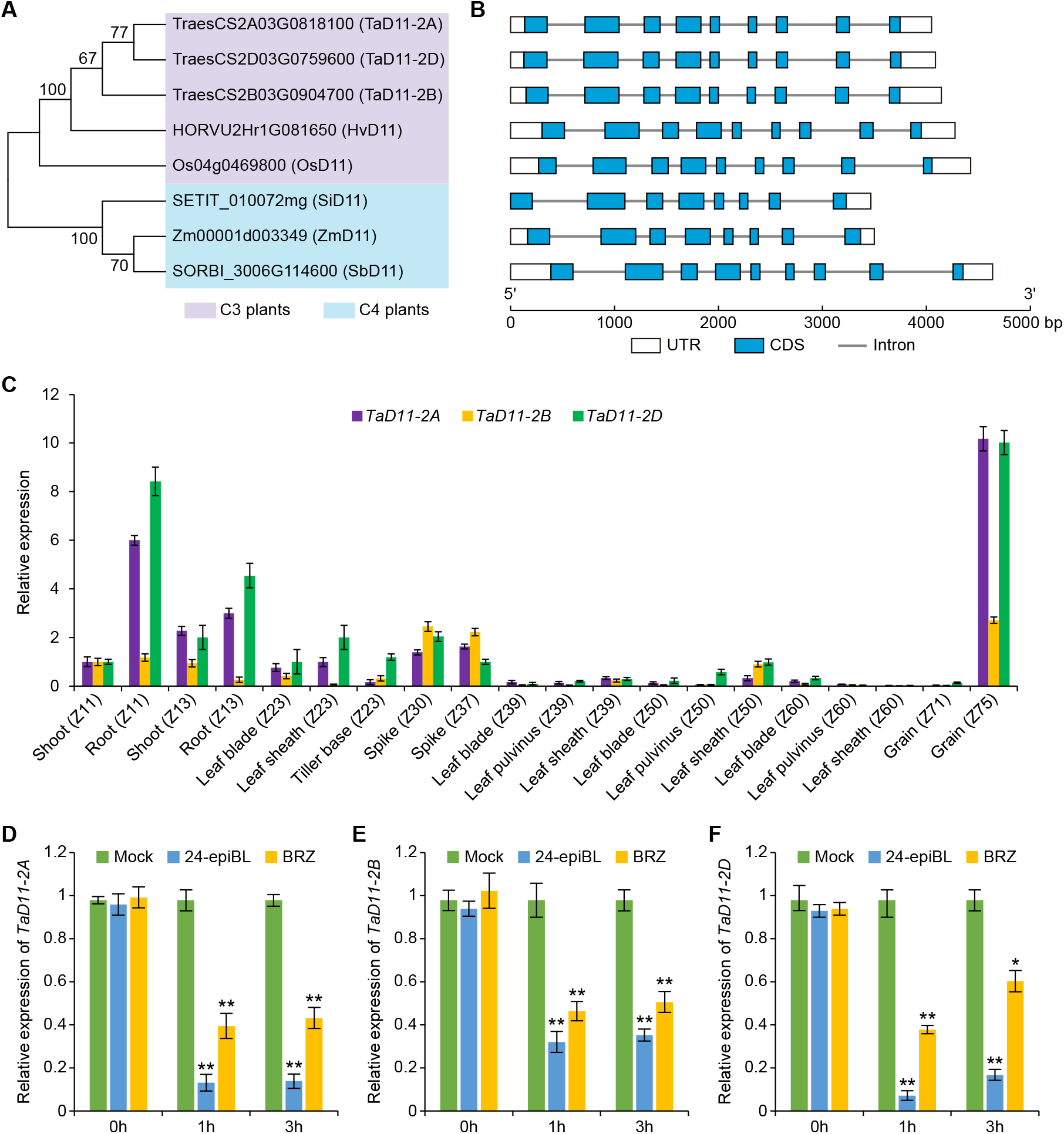
Phylogenetic relationships between D11 proteins and *TaD11* expression patterns. (A) Phylogenetic analysis of D11 protein homologs in cereals. Ta, Hv, Os, Si, Zm and Sb represent *Triticum aestivum, Hordeum vulgare, Oryza sativa, Setaria italica, Zea mays*, and *Sorghum bicolor*, respectively. A neighbor-joining tree was constructed with MEGA v10.1 and the JTT model, and was based on full-length amino acid sequences. Numbers above branches represent bootstrap support based on 1000 bootstrap replications, with a cut-off value of 50%. Purple and blue shading represents C3 and C4 plants, respectively. (B) Gene structures of D11 homologs. Untranslated region (UTR), coding sequence (CDS), and introns are represented by white boxes, blue boxes, and gray lines, respectively. Genes and their constituents are sized using the scale. (C) Expression pattern analysis of *TaD11* genes in various Chinese Spring tissue at different growth stages which followed the Zadoks scale (Zadoks *et al*., 1974). Wheat *GAPDH* was used as an internal control. Data are represented by the mean and standard deviation of three replicates. (D-F) qRT-PCR analysis of the relative expression levels of the three *TaD11* genes in Chinese Spring roots after treatment with 24-epiBL (1 _μ_M) and BRZ (1 _μ_M) after 3 hours. Values are represented by the mean and standard deviation of three independent experiments. Two-tailed Student’s *t* tests were performed between mock (distilled water) and 24-epiBL samples, and between mock and BRZ samples, respectively (\**P* < 0.05, \*\**P* < 0.01).

To clarify *TaD11* expression profiles, we investigated gene transcript levels in 20 tissues from Chinese Spring using quantitative real-time PCR (qRT-PCR) and genome-specific primers (Supplementary Table S1). All three *TaD11* genes were ubiquitously expressed in the examined tissues, with higher expression levels in root (Zadoks scale: Z11) and grain (Z75) (Fig. 1C), suggesting these genes may have potential roles in regulating root and grain development. Notably, *TaD11-2A* and *TaD11-2D* tissue expression levels were significantly higher than *TaD11-2B*. To assess *TaD11* responses to exogenous BR (24-epiBL) and the BR synthesis inhibitor, BRZ, 7-day-old Chinese Spring seedlings were treated with 1 _μ_M 24-epiBL or 1 _μ_M BRZ or a mock control (distilled water). When compared with the control, *TaD11* expression levels were all significantly decreased by 24-epiBL or BRZ treatment (Fig. 1D–F). This was not completely consistent with previously reported BR biosynthetic gene responses, i.e., *TaDWF4* genes were induced by BRZ (Hou *et al*., 2019). These observations suggested a complex feedback regulatory mechanism underpinning BR gene function in wheat.

### Functional complementarity of TaD11-2A in rice

Due to high amino acid sequence and expression pattern similarities of the three *TaD11* genes, *TaD11-2A* was selected for further functional analyses. To verify the biological function of *TaD11-2A*, we introduced an overexpression transgenic construct (p*Ubi*::*TaD11-2A*, referred to as OEW) into the rice *cpb1* mutant containing an allelic mutation of *OsD11* that exhibits clustered primary panicle branches and plant dwarfism (Wu *et al*., 2016). We showed that three independent *cpb1*-OEW lines rescued abnormal PH and panicle architecture of the *cpb1* mutant (Fig. 2A). *TaD11-2A* transcript levels in *cpb1*-OEW transgenic plants were also significantly elevated when compared with control *cpb1* plants (Fig. 2B). To further investigate if *TaD11-2A* was related to BR biosynthesis, we measured three endogenous BRs, brassinolide (BL), castasterone (CS), and 6-deoxocastasterone (6-deoxoCS) in the wild-type (WT) (CP78), *cpb1*, and transgenic plants at seeding stages. The levels of all three endogenous BRs in *cpb1*-OEW transgenic plants were significantly higher than *cpb1* and CP78 plants, especially 6-deoxoCS (Fig. 2C). Notably, when compared with CP78 plants, *cpb1*-OEW transgenic plants showed significant increases in grain length (GL) (+14.3% to +14.7%), grain area (+10.4% to +10.9%), and TGW (+13.5% to +14.6%), but a significant decrease in GW (−2.8% to −3.4%) (Fig. 2D–H). Thus *TaD11-2A* was a BR biosynthesis-related gene, with overexpression completely restoring the abnormal phenotypes of the *cpb1* mutant.

**Fig. 2.**
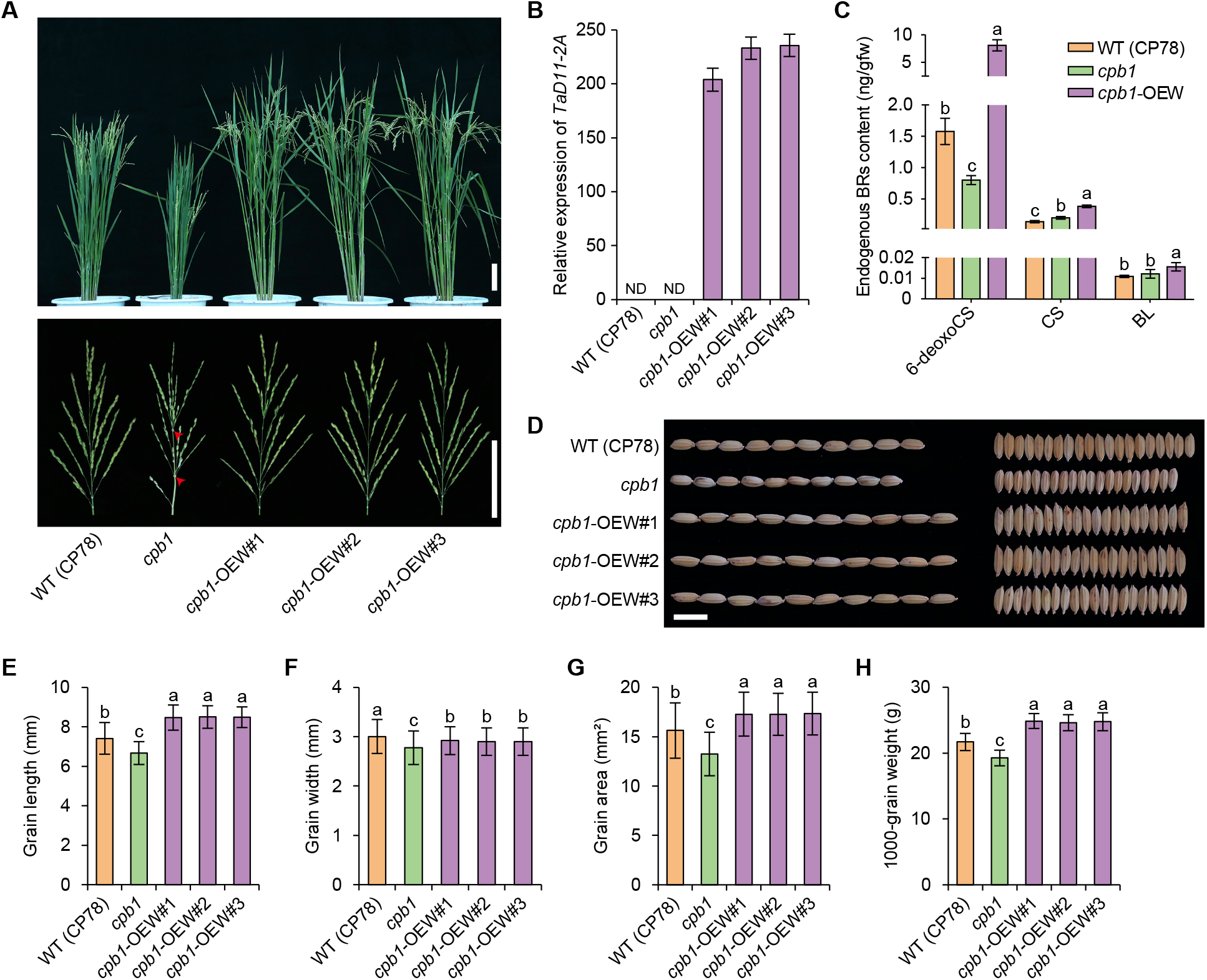
Functional complementarity validation of *TaD11-2A* in rice. (A) Comparison of plant and panicle architecture of WT (CP78), *cpb1*, and three *cpb1*-OEW transgenic lines at grain-filling stages. Scale bar = 10 cm (lower right). The red arrows indicate the position of the clustered primary branches in the *cpb1* mutant. (B) Expression analysis of *TaD11-2A* in WT (CP78), *cpb1*, and *cpb1*-OEW transgenic lines. The rice *ubiquitin* gene was used as an internal control. The expression level of *OsD11* was assigned 1.0 for transgenic lines. Values are represented as the mean and standard deviation of three independent replicates. ND = expression not detected. (C) Endogenous BR content in WT (CP78), *cpb1*, and *cpb1*-OEW in shoot tissue at the 14-day-old seeding stage. Values are mean standard deviation (*n* = 3). Gfw = grams fresh weight. (D) Comparison of grain size in WT (CP78), *cpb1*, and *cpb1*-OEW transgenic lines. Scale bar = 1 cm. (E–H) Comparison of grain length (GL) (E), grain width (GW) (F), grain area (G), and 1000-grain weight (TGW) (H) among WT (CP78), *cpb1*, and *cpb1*-OEW transgenic lines (*n* = 15). Different letters indicate significant differences in Duncan’s multiple-range tests (*P* < 0.05).

### TaD11-2A increases grain yield and grain quality in rice

To further evaluate the potential effects of *TaD11-2A* toward grain yield, the OEW construct was introduced into the *japonica* cultivar, ZH17 (Fig. 3A). When compared with the control ZH17, ZH17-OEW transgenic plants revealed dramatically elevated *TaD11-2A* expression levels and endogenous BR levels (Fig. 3B, C), significant increases in GL (+14.1% to +15.5%), GW (+2.1% to +2.8%), grain area (+20.9% to +21.1%), TGW (+13.0% to +14.7%), and GYPP (+13.7% to +14.9%) (Fig. 3D–I). We also examined total starch, amylose, and total protein levels in control ZH17 and ZH17-OEW transgenic plants to assess whether *TaD11-2A* affected rice quality traits. ZH17-OEW transgenic plants showed a significant increase in total starch levels (+3.0% to +3.4%) and a significant decrease in amylose levels (−5.0 to −5.9%), while protein levels were not significantly different (Fig. 3J–L). Therefore, *TaD11-2A* constitutive expression increased grain yield and grain quality in rice, indicating the potential of *TaD11-2A* in improving crop yield and quality traits.

**Fig. 3.**
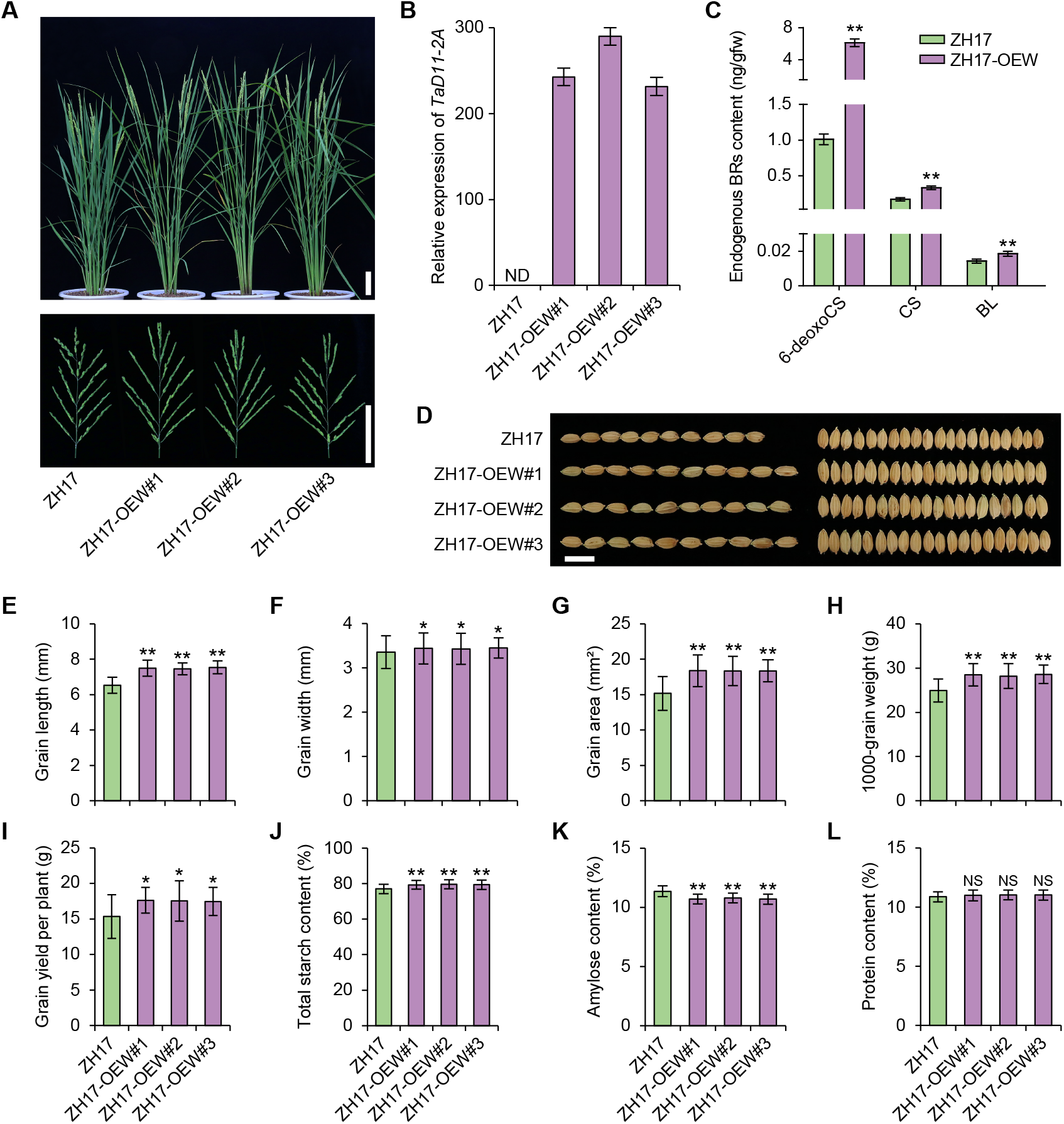
The effects of *TaD11-2A* on grain size and quality traits in rice. (A) Comparison of plant and panicle architecture between control (ZH17) and ZH17-OEW transgenic lines at the grain-filling stage. Scale bars = 10 cm (lower right). (B) Relative expression analysis of *TaD11-2A* levels between ZH17 and ZH17-OEW at the grain-filling stage. Rice *ubiquitin* was used as an internal control. The data are represented by the mean and standard deviation of three independent replicates. The expression level of *OsD11* was assigned 1.0 for the transgenic lines. ND = expression not detected. (C) Endogenous BR contents between ZH17 and ZH17-OEW plants in shoot tissue at the 14-day-old seeding stage. Values are mean standard deviation (*n* = 3). Gfw = gram fresh weight. (D) Comparison of grain size between the ZH17 control and ZH17-OEW transgenic lines. Scale bar = 1 cm. (E–L) Comparison of grain length (GL) (E), grain width (GW) (F), grain area (G), 1000-grain weight (TGW) (H), grain yield per plant (GYPP) (I), total starch content (J), amylose content (K), and protein content (L) between ZH17 and ZH17-OEW transgenic lines (for yield-related traits, *n* = 15; for quality-related traits, *n* = 3). Two-tailed Student’s *t* tests were performed between ZH17 and ZH17-OEW (NS = not significant; \**P* < 0.05, \*\**P* < 0.01).

### Phenotypic characterization of the wheat tad11-2a mutant

To investigate the role of *TaD11* in wheat, we screened the *Triticum turgidum* cv. Kronos TILLING population (Krasileva *et al*., 2017) and identified the *tad11-2a* mutant (Kronos2702). After three generations of backcrossing, the homozygous *tad11-2a* mutant line was selected for further analysis (Fig. 4A, B). Our qRT-PCR data revealed that *TaD11-2A* expression in grain (Z75) was significantly lower in the mutant than the WT (Kronos), while no significant differences were identified in other tissues (Fig. 4C). Agronomic trait investigations showed that when compared with the WT (Kronos), the *tad11-2a* mutant exhibited significant decreases in PH (−18.9%), spike length (SL) (−12.5%), spikelet number per spike (SNPS) (−5.5%) and grain number per spike (GNPS) (−31.0%) (Fig. 4D, F–I). Furthermore, GL, GW, and the grain area of the *tad11-2a* mutant were significantly reduced by 5.6%, 8.0%, and 11.4%, respectively, when compared with WT (Kronos) (Fig. 4E, J–L), leading to serious decreases in TGW (−31.0%) and GYPP (−41.4%) (Fig. 4M, N). We also tested three quality related traits in the mutant and WT and found that protein and wet gluten content (WGC) levels were significantly higher in the mutant than the WT by 17.1% and 14.2% (Fig. 3O, Q), respectively, while no significant difference was observed for test weight (Fig. 4P). In addition, the wheat lamina joint inclination assay showed that the *tad11-2a* mutant was more sensitive to 24-epiBL treatment than the WT (Fig. 4R). Thus, *TaD11-2A* was a BR-related gene affecting grain yield and grain quality in wheat.

**Fig. 4.**
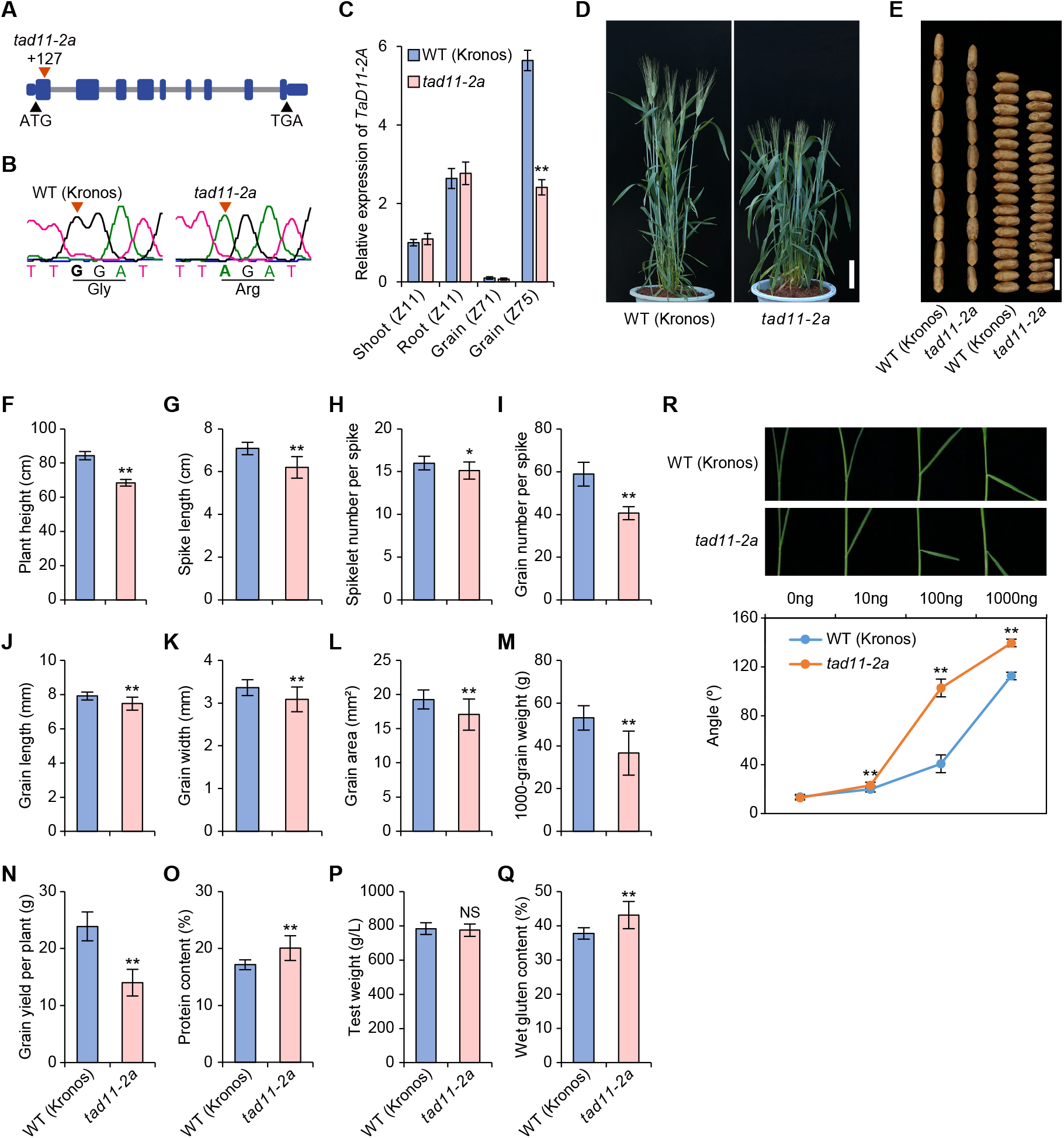
Morphological characters of the *tad11-2a* mutant. (A) Genomic structure of *TaD11-2A*. The red triangle is the variant position in the *tad11-2a* mutant. (B) Sequencing validation of the *tad11-2a* mutation site. (C) Expression analysis of *TaD11-2A* in WT (Kronos) and the *tad11-2a* mutant. Wheat *GAPDH* was used as an internal control. The data are the mean plus the standard deviation of three replicates. (D) Plant architecture of the WT (Kronos) and *tad11-2a* mutant at the heading stage. Scale bar = 10 cm. (E) Grain shape of WT (Kronos) and the *tad11-2a* mutant. Scale bar = 1 cm. (F–Q) Comparison of 12 agronomic traits, including, plant height (PH) (F), spike length (SL) (G), spikelet number per spike (SNPS) (H), grain number per spike (GNPS) (I), grain length (GL) (J), grain width (GW) (K), grain area (L), 1000-grain weight (TGW) (M), grain yield per plant (GYPP) (N), protein content (PC) (O), test weight (TW) (P), and wet gluten content (WGC) (Q) in the WT (Kronos) and *tad11-2a* mutant (*n* = 15). (R) Comparison of lamina inclination in the WT (Kronos) and *tad11-2a* mutant after 24-epiBL treatment (*n* = 15). Values are the mean plus standard deviation. Two-tailed Student’s *t* tests were performed between the WT (Kronos) and *tad11-2a* mutant (NS = not significant; \**P* < 0.05, \*\**P* < 0.01).

### Haplotype identification and molecular marker development of TaD11

To explore the relationship between natural variations in *TaD11* and agronomic traits, we analyzed polymorphisms in the coding and 2-kb promoter regions of *TaD11* in 60 wheat accessions with high genetic diversity (Supplementary Table S3) from 17 countries. Two *TaD11-2A* haplotypes were characterized by nine single nucleotide polymorphisms (SNPs) and two insertions/deletions (InDels), most of which were located in promoter or intron regions, except InDel2 in the 5’ untranslated region (UTR) (Supplementary Fig. S2A). An InDel marker (DA-InDel2) was developed to distinguish the two *TaD11-2A* haplotypes by denaturing polyacrylamide gel (Supplementary Fig. S2A). For *TaD11-2B*, ten SNPs in promoter or intron regions were identified and resulted in two haplotypes (Supplementary Fig. S2B). A derived cleaved amplified polymorphic sequence (dCAPS) marker, DB-SNP10 was designed to detect the two *TaD11-2B* haplotypes by agarose gel (Supplementary Fig. S2B). For *TaD11-2D*, no nucleotide variations were observed in promoter and coding regions (Supplementary Fig. S2C).

### TaD11 haplotypes are associated with grain yield and grain quality in wheat

We scanned 314 wheat accessions (Supplementary Table S4) from 23 countries using the Wheat 55K SNP array, and filtered it to generate 24838 high-quality unique SNPs. Population structure analysis showed that the natural population was divided into four subgroups (Fig. 5A–C). Based on a total of 24840 markers from *TaD11* haplotypes and the Wheat 55K SNP array in the natural population, and combined with 15 agronomic traits in seven environments during the years 2017–2020 (Supplementary Table S5), we performed a genome-wide association analysis (GWAS) using the general linear model (GLM) and principal component analysis (PCA). *TaD11-2A* was significantly associated with ten yield-related traits (*P* < 0.05), such as PH (seven environments), awn length (AL) (seven environments), and GYPP (four environments) (Fig. 5D; Supplementary Table S6), and also with two quality-related traits, protein content (PC) (one environment) and WGC (two environments) (Supplementary Fig. S3A; Supplementary Table S6). In contrast, *TaD11-2B* was significantly associated with only three yield-related traits, spike number (SN) (two environments), SNPS (one environment), and TGW (one environment) (Supplementary Fig. S4A; Supplementary Table S6), and with two quality traits, protein (two environments) and WGC (one environment) (Supplementary Fig. S3B; Supplementary Table S6), which may have been due to low minor allele frequency (MAF = 2.5%). We also performed linkage disequilibrium (LD) analysis on 5-Mb regions upstream and downstream of *TaD11* genes, and found that *TaD11-2A* exhibited weak LD (*r*^2^ < 0.5) (Fig. 5E), while *TaD11-2B* and *TaD11-2D* regions had no LD due to few markers. These data indicated the homolog-specific functions of *TaD11*.

**Fig. 5.**
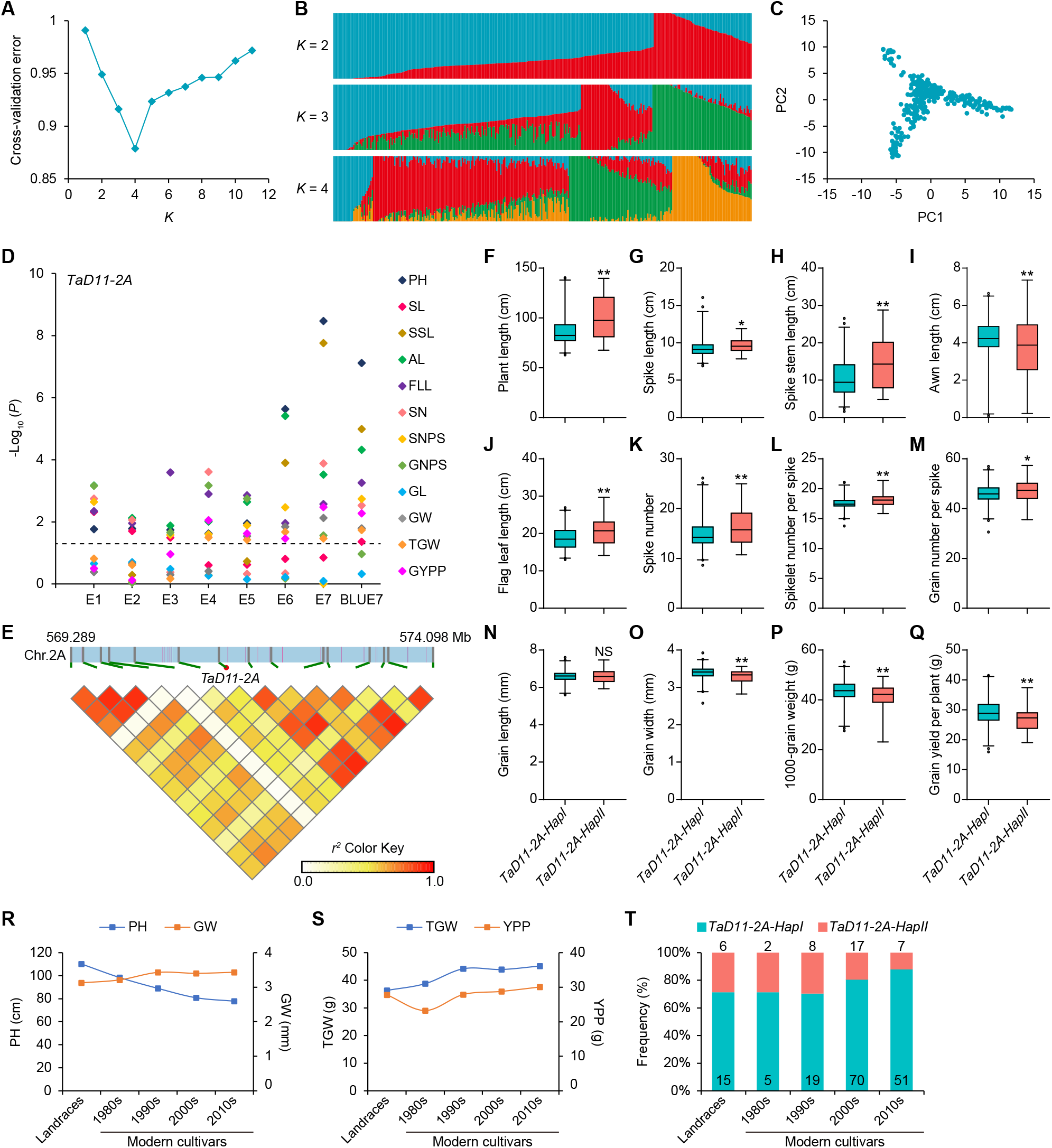
Haplotype analysis of *TaD11-2A* associations with yield-related traits in wheat. (A–C) Population structure analyses were performed on 314 wheat accessions using cross-validation error (A), stacked bar plots of ancestry relationship (B), and principal component analyses (C). (D) Genome-wide association analysis of *TaD11-2A* with 12 yield-related traits (PH, plant height; SL, spike length; SSL, spike stem length; AL, awn length; FLL, flag leaf length; SN, spike number; SNPS, spikelet number per spike; GNPS, grain number per spike; GL, grain length; GW, grain width; TGW, 1000-grain weight; GYPP, grain yield per plant). E1–E7 represents association analysis in seven different environments. BLUE7 indicates association analysis based on the best linear unbiased estimate (BLUE) values calculated from seven data environments. Negative log_10_-transformed *P* values are plotted. A black horizontal dotted line indicates the threshold value for significant associations (*P* < 0.05). (E) Linkage disequilibrium (LD) analysis spanning the physical position from 569.289 to 574.098 Mb (IWGSC RefSeq v2.1) of chromosome 2A. The color key indicates the level of LD (*r*^2^) between variants. (F–Q) Comparison of 12 yield-related traits, including PH (F), SL (G), SSL (H), AL (I), FLL (J), SN (K), SNPS (L), GNPS (M), GL (N), GW (O), TGW (P), and GYPP (Q) between *TaD11-2A* haplotypes based on BLUE values. Two-tailed Student’s *t* tests were performed between *TaD11-2A* haplotypes (NS = not significant; \**P* < 0.05, \*\**P* < 0.01). (R, S) Changes in PH and GW (R) and TGW and GYPP (S) in 200 Chinese wheat accessions over several decades. In this population, there were 21 landraces, and 7, 27, 87, and 58 modern cultivars bred in the 1980s, 1990s, 2000s, and 2010s, respectively. Data for PH, GW, TGW, and GYPP are represented by a line chart using BLUE values. (T) Changes in the frequency of *TaD11-2A* haplotypes across the history of wheat breeding in China.

To further clarify the effects of *TaD11* haplotypes on agronomic traits, we calculated best linear unbiased estimate (BLUE) values for 15 traits in 314 wheat accessions using phenotypic data in multiple environments. Overall, *TaD11-2A-HapI* was more favorable when compared to *TaD11-2A-HapII*, as evidenced by significant decreases in PH (−12.7%), spike stem length (SSL) (−25.9%) and flag leaf length (FLL) (−10.1%), while significant increases were observed in GW (+3.3%), TGW (+5.0%) and GYPP (+7.6%), despite decreases in GNPS (−2.6%) and PC (−3.9%) (Fig. 5F–Q; Supplementary Fig. S3B-D). Furthermore, *TaD11-2B-HapII* was a rare variant, and when compared with *TaD11-2B-HapI*, no significant changes were observed in all traits except for FLL, which were significantly shorter (−20.5%) (Supplementary Fig. S3F–H; Supplementary Fig. S4B–M). These findings suggested *TaD11-2A-HapI* was a novel marker for grain yield, with potential use in wheat breeding programs.

### TaD11-2A-HapI is positively selected in wheat breeding

To determine the selection characteristics of *TaD11* haplotypes in wheat breeding, we assessed the frequency variations of *TaD11-2A* and *TaD11-2B* haplotypes in 200 Chinese wheat accessions from different breeding years (Supplementary Table S4). According to BLUE data from these accessions, PH showed a gradual decrease from landraces to modern cultivars, while GW, TGW, and GYPP showed a gradual increase (Fig. 5R, S). Correspondingly, the frequency of *TaD11-2A-HapI* increased from 71.43% for landraces to 87.93% for modern cultivars of the 2010s (Fig. 5T), while the *TaD11-2B-HapI* frequency remained essentially unchanged (Supplementary Fig. S4N). These results suggested that *TaD11-2A-HapI* underwent positive selection in wheat breeding processes in China.

We also examined the global geographic distributions of *TaD11-2A* and *TaD11-2B* haplotypes in 314 wheat accessions from six continents (Supplementary Table S4). For *TaD11-2A, HapI* had the highest overall global proportion, especially in Asia and North America, where proportions were 80.40% and 95.24%, respectively, while other continents had a lower proportion of *HapI* when compared to *HapII*, probably due to the small number of wheat accessions (Supplementary Fig. S5A). For *TaD11-2B*, the overall percentage of *HapI* was 96.76% globally, while *HapII* was only marginally distributed in Asia and Oceania (Supplementary Fig. S5B). These results indicated that *TaD11-2A-HapI* had been subject to positive global wheat breeding selection.

### Natural variations of OsD11 affect panicle length and GW in rice

Previous studies revealed that *OsD11* regulated multiple agronomic traits, such as PH, panicle architecture, and grain size (Tanabe *et al*., 2005; Wu *et al*., 2016), but potential associations between its natural genetic variations and yield-related traits in rice have remained unclear. Therefore, we investigated polymorphisms in the coding and 2-kb promoter regions of *OsD11* in 3024 germplasm accessions based on sequence variation data of the 3K rice genome panel (Alexandrov *et al*., 2015). We identified 42 polymorphic sites and performed association analyses with four yield-related traits, including panicle length (PL), GL, GW, and TGW. Several sites were significantly associated with yield-related traits (*P* < 0.05), especially the SNP13 on intron 1 of *OsD11* which had the highest association with PL and GW, as well as a significant association with GL and TGW (Fig. 6A, B; Supplementary Table S7). The LD of this SNP with other polymorphic sites was not significant (*r*^2^ < 0.1) (Fig. 6C). Further analysis indicated this SNP was divided into two haplotypes, *OsD11-HapI* (REF: T) and *OsD11-HapII* (ALT: C). When compared with *OsD11-HapI, OsD11-HapII* showed a significant decrease in PL (−12.5%) and a significant increase in GW (+10.1%) (Fig. 6D, E). In addition, the majority of *xian/indica* (*XI*) and *AUS* accessions contained *OsD11-HapI*, while 12.4% of *geng/japonica* (*GJ*) accessions carried *OsD11-HapII*, including 27.2% in *temperate japonica* (*GJ_tmp*), 0.8% in *tropical japonica* (*GJ_trp*), 13.3% in *subtropical japonica* (*GJ_subtrp*), and 12.2% in *admix* (*GJ_adm*) (Fig. 6F). Taken together, *OsD11-HapII* was a rare allelic variant of *OsD11* and was mainly distributed in the temperate regions of Asia.

**Fig. 6.**
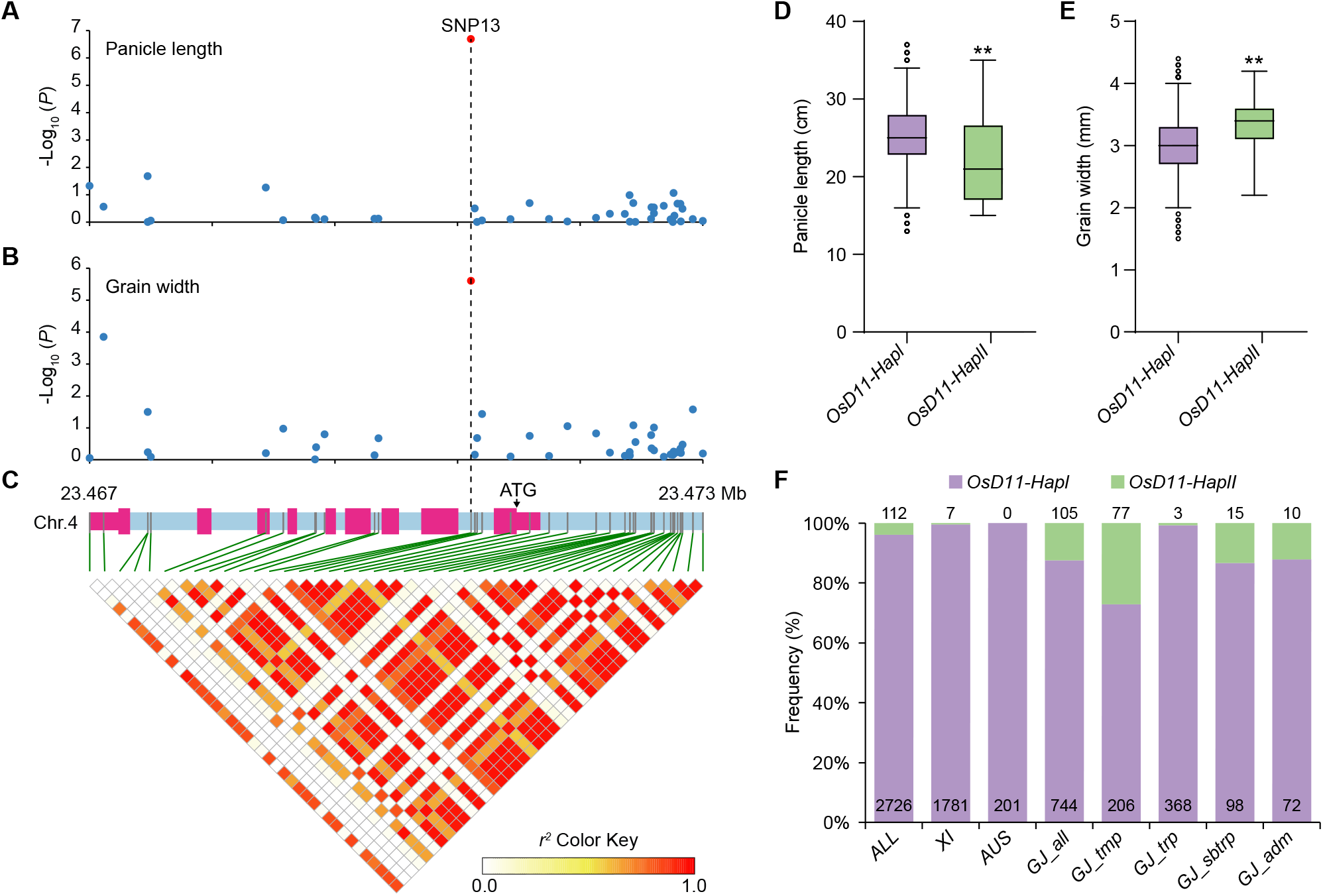
Haplotype analysis of *OsD11* associations with panicle length and grain width in rice. (A, B) Association analysis of *OsD11* with panicle length (PL) and grain width (GW) in the 3K rice genome panel. Red dots represent the most significantly associated polymorphic sites in *OsD11* with PL and GW, respectively. (C) Triangle matrix of pairwise linkage disequilibrium (LD) of DNA polymorphisms in the coding and 2-kb promoter regions of *OsD11*. The color key indicates the level of LD (*r*^2^) between variants. (D, E) Phenotypic comparison of *OsD11* haplotypes with PL and GW. Two-tailed Student’s *t* tests were performed between haplotypes (\*\**P* < 0.01). (F) The frequency distribution of *OsD11* haplotypes in different subgroups; *XI, xian/indica*; *GJ_all, geng/japonica*; *GJ_tmp, temperate japonica*; *GJ_trp, tropical japonica*; *GJ_sbtrp, subtropical japonica*; and *GJ_adm, admix*.

### TaD11-2A influences root length and salt tolerance in rice and wheat

Given the high *TaD11* expression in wheat roots (Fig. 1C), we hypothesized these genes may have important roles in the regulation of root development and abiotic stress tolerance. Thus, comparative root phenotype analyses and salt and drought tolerance studies were conducted at seeding stages in rice and wheat, respectively. In rice, ZH17-OEW transgenic plants overexpressing *TaD11-2A* had significantly increased shoot height (+12.3% to +13.9%) and significantly shorter total root length (−28.2% to −30.2%) when compared with control plants, ZH17 (Fig. 7A). Our salt and drought tolerance assays revealed that the survival rates of ZH17-OEW transgenic plants under salt stress (1% NaCl) were significantly higher than controls (ZH17). In contrast, the survival rates of ZH17-OEW transgenic plants under drought stress (20% PEG6000) were significantly lower than controls (Fig. 7B, C). In wheat, shoot height and total root length of the *tad11-2a* mutant were both significantly decreased by 26.2% and 22.4%, respectively, when compared with the WT (Kronos) (Fig. 7D). Under salt stress (1.5% NaCl), the survival rate of the *tad11-2a* mutant was significantly reduced compared to the WT (Kronos), whereas under drought stress (20% PEG6000), no significant differences were observed between survival rates of test and control plants (Fig. 7E–F).

**Fig. 7.**
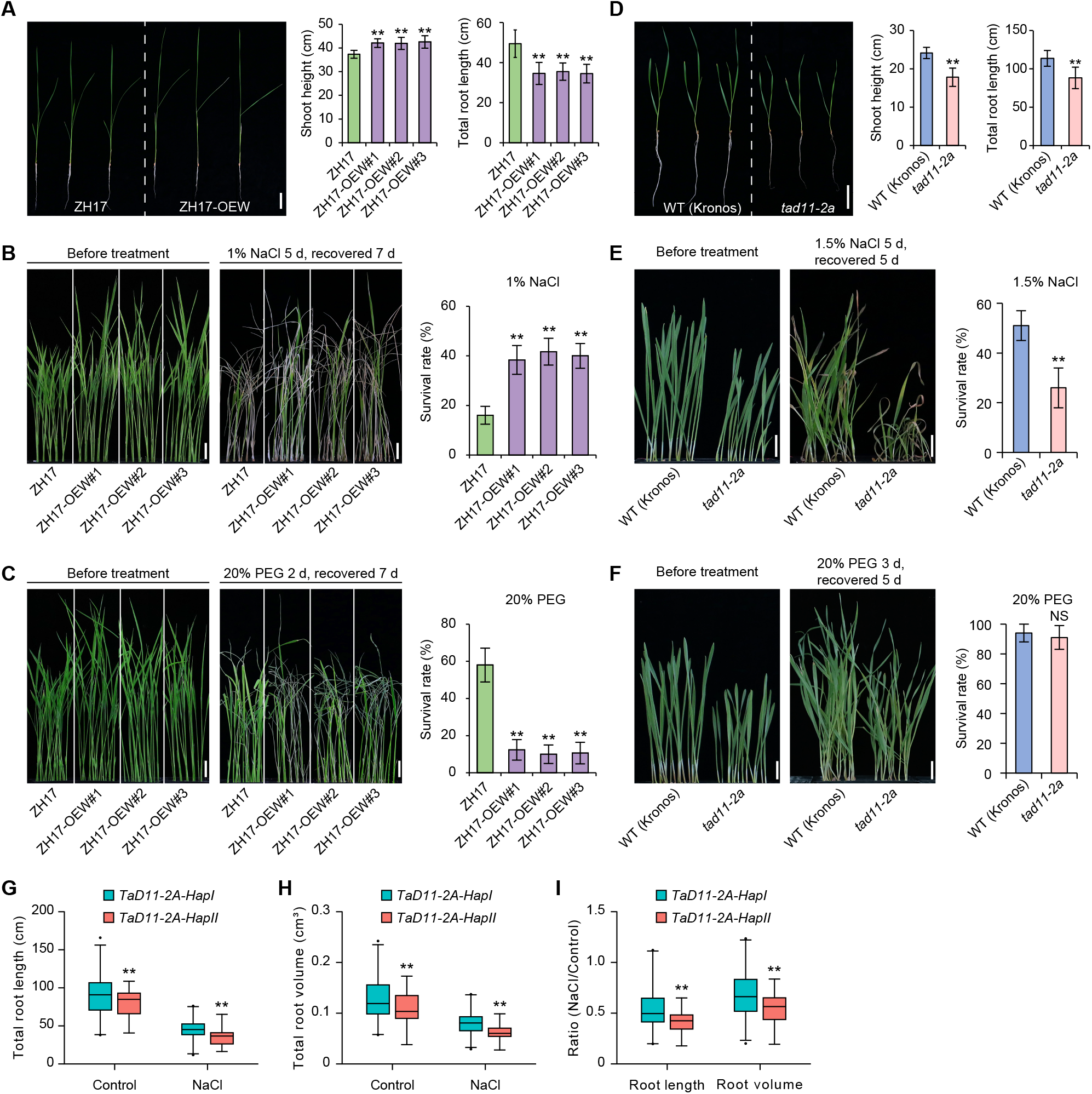
*TaD11-2A* affects root length and salt tolerance at seeding stages in rice and wheat. (A) Comparison of shoot height and root length between control (ZH17) and ZH17-OEW transgenic plants (*n* = 20) at seeding stages in rice. Scale bar = 5 cm. (B) Phenotypes and survival rates (*n* = 6) of ZH17 and ZH17-OEW plants during salt stress (1% NaCl). Scale bar = 2 cm. (C) Phenotypes and survival rates (*n* = 6) of ZH17 and ZH17-OEW plants during drought stress (20% PEG6000). Scale bar = 2 cm. (D) Comparison of seeding height and root length between the WT (Kronos) and *tad11-2a* mutant (*n* = 20) at seeding stages in wheat. Scale bar = 5 cm. (E) Phenotypes and survival rates (*n* = 6) of WT (Kronos) and *tad11-2a* plants during salt stress (1.5% NaCl). Scale bar = 2 cm. (F) Phenotypes and survival rates (*n* = 6) of WT (Kronos) and *tad11-2a* plants during drought stress (20% PEG6000). Scale bar = 2 cm. (G–I) Comparison of total root length (G), total root volume (H), and ratio (NaCl/control) (I) of wheat accessions (E3) in the natural population at the seedling stage under normal conditions and salt treatment (*n* = 6). Values are mean standard deviation. Two-tailed Student’s *t* tests were performed between ZH17 and ZH17-OEW plants, WT (Kronos) and *tad11-2a* plants, and *TaD11-2A* haplotypes, respectively (NS = not significant; \*\**P* < 0.01).

To clarify the effects of natural variation in *TaD11* on root length, root phenotyping was performed on wheat accessions (E3) in the natural population. *TaD11-2A-HapI* had longer total root length (+11.4% to +19.8%) and larger total root volume (+15.3% to +25.4%) relative to *TaD11-2A-HapII* under both normal and salt treatment conditions (Fig. 7G, H). Interestingly, the total root length ratio and total root volume ratio (NaCl/control) for *TaD11-2A-HapI* was significantly higher than *TaD11-2A-HapII* under salt treatment (Fig. 7I), suggesting a greater salt tolerance. Thus, *TaD11-2A* had an important role regulating root development and salt tolerance at the wheat seeding stage.

## Discussion

BRs represent the sixth class of phytohormones, and while the major biosynthesis pathways have been largely elucidated, it is unclear how to finely control BR-related genes and regulate BR bioactivity levels in response to various developmental processes and environmental stresses (Bajguz *et al*., 2020; Zhao and Li, 2012). In rice, *D11* is a key gene in the BR biosynthetic pathway, with several of its alleles involved in the regulation of a variety of agronomic traits, including PH, leaf angle, panicle architecture, grain shape, and grain number (Tanabe *et al*., 2005; Wu *et al*., 2016; Zhou *et al*., 2017). *OsDWARF4* and *D11* are functionally redundant, and mutation of *OsDWARF4* alone exerts limited effects on BR biosynthesis and plant morphology (Sakamoto *et al*., 2006). In our study, we identified *TaD11* genes in wheat; the *tad11-2a* mutant exhibited a typical BR deficient phenotype characterized by dwarfism and small round seeds, shorter SL, reduced grain number, and sensitivity to 24-epiBL (Fig. 4). *TaD11-2A* overexpression rescued the defective phenotypes in the rice *cpb1* mutant and increased endogenous BR levels (Fig. 2), suggesting *TaD11-2A* had conserved functions. The *TaD11* genes were significantly down-regulated during 24-epiBL or BRZ treatment (Fig. 1D-F), which was not entirely consistent with BR responses in wheat; *TaDWF4* genes were induced by BRZ treatment (Hou *et al*., 2019). BRZ is a triazole compound and specifically inhibits BR biosynthesis by blocking the cytochrome P450 steroid, C-22 hydoxylase which is encoded by *AtDWF4* (Asami *et al*., 2000; Asami *et al*., 2001). Thus, *TaD11* and *TaDWF4* may have different roles in the BR biosynthetic pathway in wheat, leading to different responses to BRZ treatment.

Grain yield is the most important agronomic trait in grain crops; extensive efforts have been made to improve outputs, especially for the three major global cereal crops, rice, maize, and wheat (Vriet *et al*., 2012). Engineering BR biosynthesis and signaling pathways may provide effective strategies for improving crop production. For example, the specific up-regulation of the BR biosynthesis gene, *OsD11* increased grain size and GYPP in rice (Wu *et al*., 2016), whereas editing the BR signal transduction gene, *ZmRAVL1* enhanced high-density yields in maize (Tian *et al*., 2019). However, the potential exploitation of natural variations in BR-related genes toward wheat grain yield is rarely reported. In this study, the constitutive overexpression of *TaD11-2A* in rice significantly improved grain size, grain weight, GYPP, and grain quality, but was unsuitable for dense planting due to loose plant architecture and excessive leaf angles (Fig. 3). These factors suggested that controlling the precise and specific expression of *TaD11* in grain development was essential for further yield improvements. Our analyses on *TaD11* genetic variability in the natural population provided evidence that a specific haplotype, *TaD11-2A-HapI*, was positively associated with GW, TGW, and GYPP, and was positively selected in wheat breeding (Fig. 6O–T). Interestingly, natural variations in *OsD11* were significantly associated with several yield-related traits in rice, including PL and GW, suggesting *D11* may have conserved and specific roles in different crops. Taken together, *TaD11-2A-HapI* represented an effective molecular marker for wheat breeding programs.

Roots are important determinants of crop yield (Vriet *et al*., 2012). Root growth is promoted by low concentrations and inhibited by high concentrations of exogenous BR (Mussig *et al*., 2003). Many BR-deficient mutants have reduced root length and lateral roots, indicating the positive effect of BRs on root development at physiological concentrations (Bao *et al*., 2004; Ferguson *et al*., 2005). In wheat, BR has a conserved function in regulating root length, and also novel roles by controlling lateral root emergence and root diameter (Hou *et al*., 2019). In our study, endogenous BR levels were significantly enhanced in ZH17-OEW transgenic plants, which also exhibited increased shoot height and shorter root length at seedling stages (Fig. 7A), suggesting shoots and roots may have different physiological BR concentrations. When compared with control ZH17 plants, ZH17-OEW transgenic plants also showed enhanced salt tolerance and decreased drought resistance (Fig. 7B, C), suggesting BR physiological responses due to different abiotic stresses may be different. In addition, the *tad11-2a* mutant exhibited reduced root length and decreased salt tolerance (Fig. 7D, E), and *TaD11-2A-HapI* showed longer total root length and larger total root volume under normal and salt stress conditions (Fig. 7G, H). Notably, *TaD11-2A-HapI* had a higher total root length ratio and total root volume ratio (NaCl/control) under salt stress (Fig. 8I) conditions. Thus, *TaD11-2A* has important roles regulating root development in wheat, and its favorable haplotype may have increased salt tolerance.

## Supplementary data

**Fig. S1**. Multiple sequence alignment of TaD11 and homologs.

**Fig. S2**. Haplotypes and molecular markers of *TaD11*.

**Fig. S3**. Haplotype analysis of *TaD11* associations with quality-related traits in wheat. **Fig. S4**. Haplotype analysis of *TaD11-2B* associations with yield-related traits in wheat.

**Fig. S5**. Global distribution of *TaD11* haplotypes.

**Table S1**. Primers used in this study.

**Table S2**. TaD11 information and homologs in cereals.

**Table S3**. Sixty wheat accessions for polymorphism analysis.

**Table S4**. Origins and *TaD11* haplotypes of 314 wheat accessions.

**Table S5**. Information on the seven environments used in this study.

**Table S6**. *P* values for association analyses between *TaD11* haplotypes and 15 agronomic traits in seven environments.

**Table S7**. *P* values for association analyses between *OsD11* natural variations and four yield-related traits in rice.

## Acknowledgments

This work was supported by the National Natural Science Foundation of China (Grant No. 32072051), Natural Science Foundation of Shandong Province, China (Grant No. ZR2019PC003), the Major Basic Research Project of Natural Science Foundation of Shandong Province, China (Grant No. ZR2019ZD16), the Agricultural Variety Improvement Project of Shandong Province (Grant No. 2021LZGC013), the Youth Innovation Technology Support Planning Project for Institution of Higher Education of Shandong Province, China (Grant No. 2019KJF002), the National Natural Science Foundation for Young Scholars of China (Grant No. 32101726), and the Yantai

Institute of Replacing Old Growth Drivers with New Ones & Yantai Demonstration Base for the Transfer and Transformation of Sci-tech Achievements Funded Project (2019XJDN007). We thank Prof. Lubin Tan (China Agricultural University) for sharing the *cpb1* mutant.

## Author contributions

YW and FC designed the experiments and managed the project. HX, HS, JD, CM, JL, ZL, YW, JJ, XH, and MW performed experiments; CZ, RQ, JW, and FN supervised the project and provided technical assistance. YW, HX, and HS analyzed data and wrote the manuscript. All authors read and agreed to the final published version of the manuscript.

## Data availability

All data related to this manuscript can be found within this paper and its Supplementary data.

