## Supplementary Figures S1-S5 for "The brassinosteroid biosynthesis gene *TaD11-2A* controls grain size and its elite haplotype improves wheat grain yields"

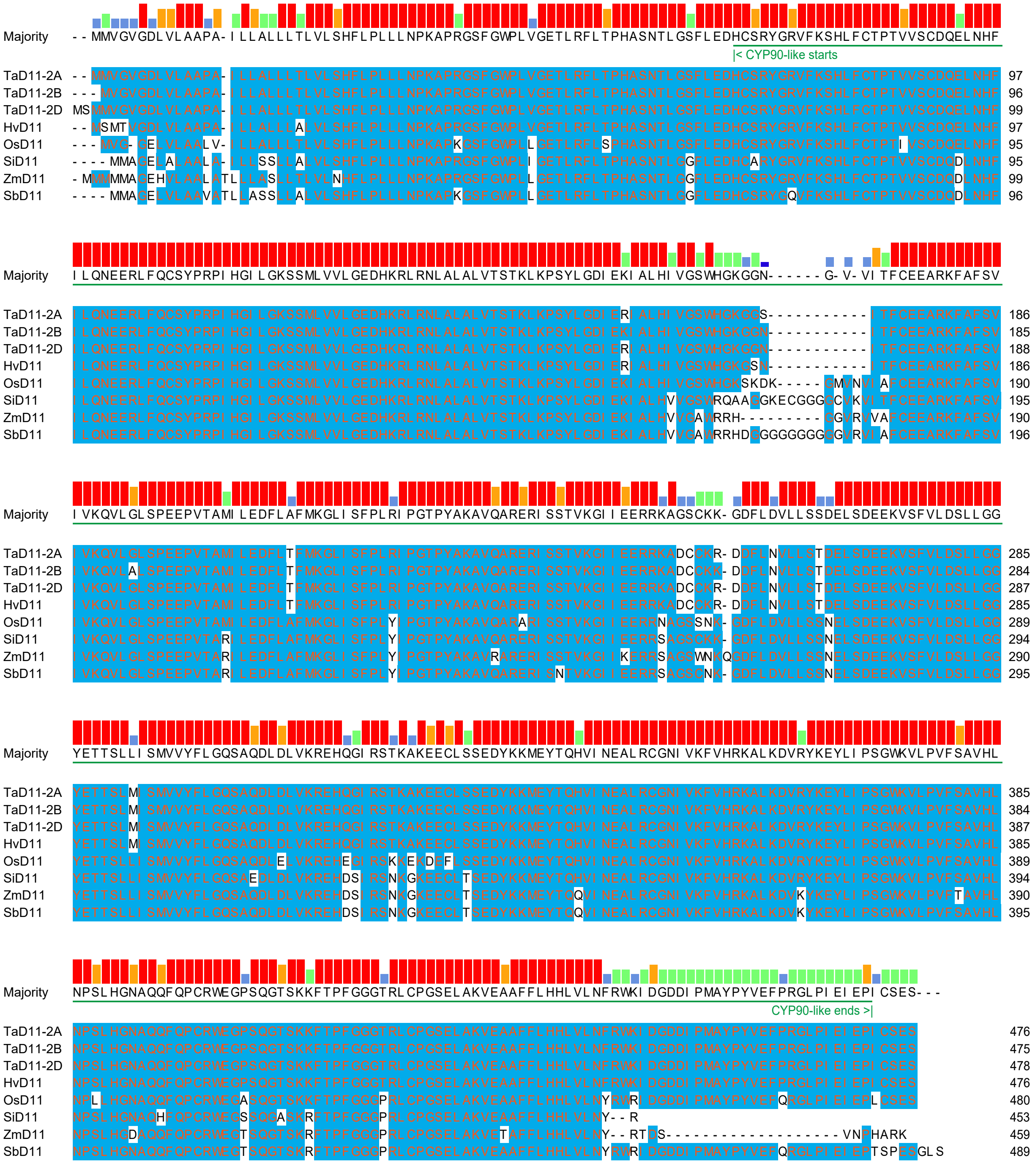


**Fig. S1.** Multiple sequence alignment of TaD11 and homologs. This alignment graphic was generated by MegAlign of DNASTAR using the ClustalW program for multiple protein sequence alignments. The major green line refers to the CYP90-like domain.

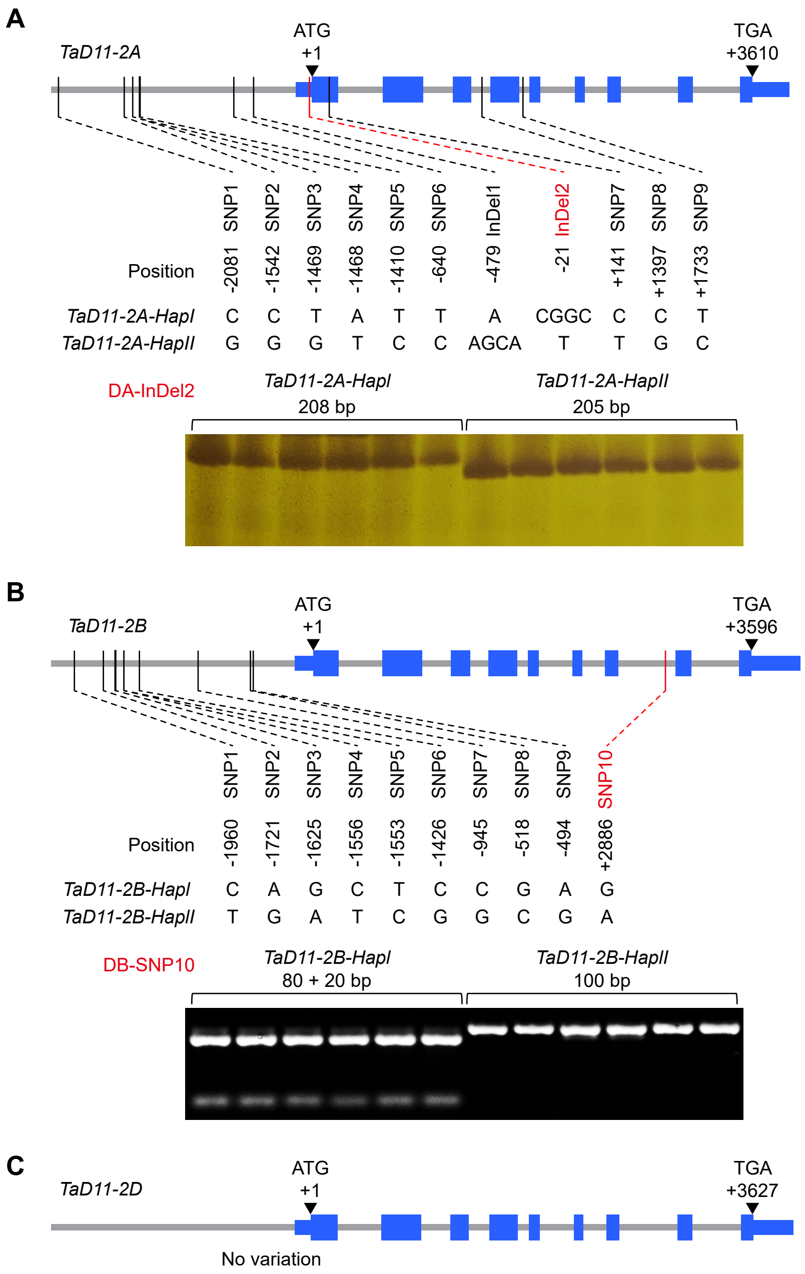


**Fig. S2.** Haplotypes and molecular markers of *TaD11*. (A) Two haplotypes and molecular marker of *TaD11-2A*. An InDel marker was developed based on the InDel2. (B) Two haplotypes and molecular marker of *TaD11-2B*. A derived cleaved amplified polymorphic sequence (dCAPS) marker was designed based on the digest of SNP10 with the restriction endonuclease, *AluI*. (C) Nucleotide variation in *TaD11-2D*. The ATG start codon is designated as position 1 bp. Exons are indicated by blue rectangles; promoters and introns are indicated by gray rectangles. Variations and relative positions are shown below the *TaD11* gene structures.

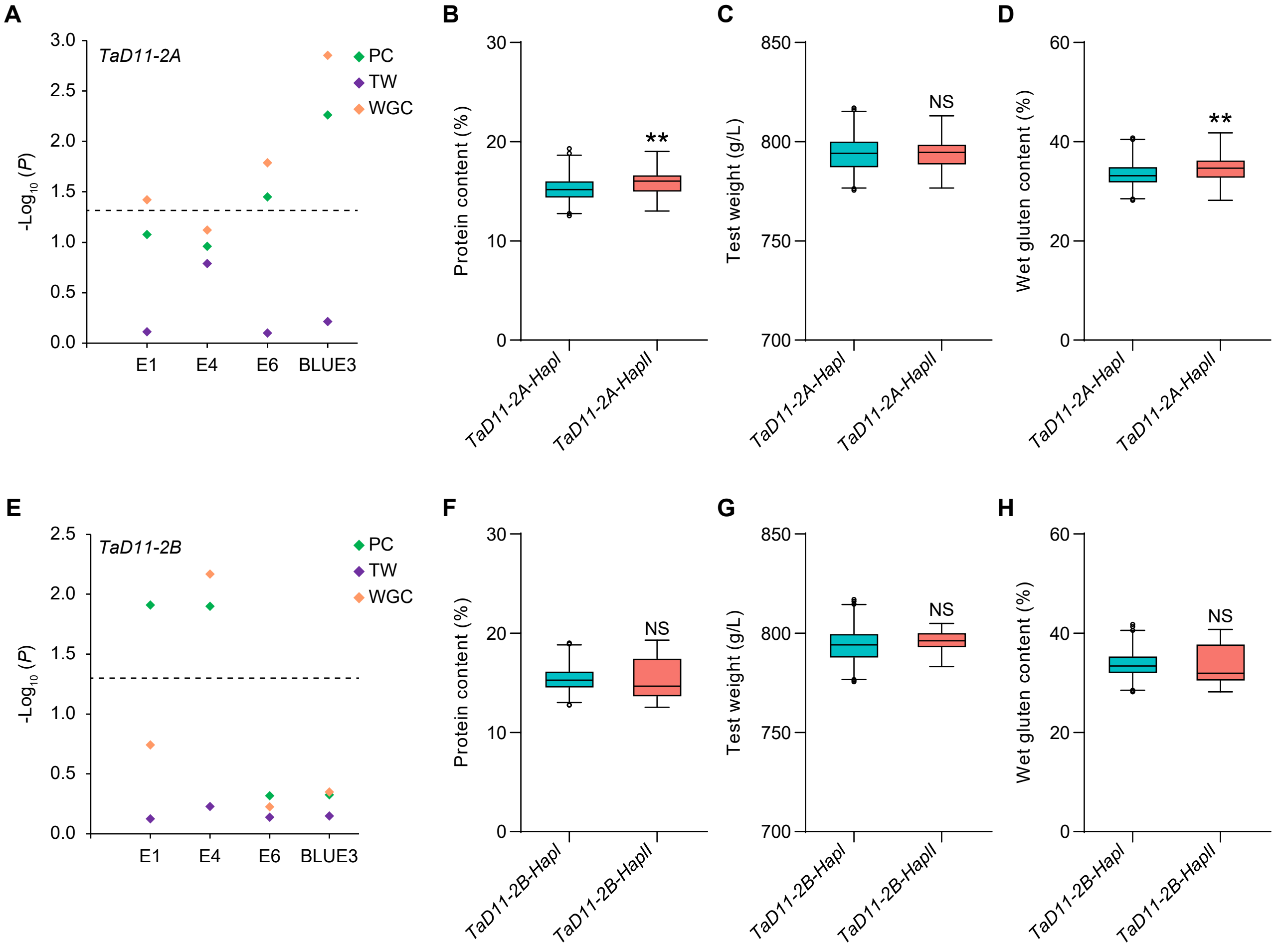


**Fig. S3.** Haplotype analysis of *TaD11* associations with quality-related traits in wheat. (A) Genome-wide association analysis of *TaD11-2A* with three quality-related traits. PC, protein content; TW, test weight; and WGC, wet gluten content. E1, E4, and E6 indicate association analysis in three different environments. BLUE3 indicates association analysis based on the best linear unbiased estimate (BLUE) values calculated from three environment data (E1, E4 and E6). Negative $\log_{10}$-transformed $P$ values are plotted. A black horizontal dotted line indicates the threshold value for significant associations ($P < 0.05$). (B–D) Comparison of three quality-related traits, including PC (B), TW (C), and WGC (D), between *TaD11-2A* haplotypes based on BLUE values. (E) Genome-wide association analysis of *TaD11-2B* with three quality-related traits. (F–H) Comparison of three quality-related traits, including PC (F), TW (G), and WGC (H) between *TaD11-2B* haplotypes based on BLUE values. Two-tailed Student's $t$ tests were performed between haplotypes (NS = not significant; $^{**}P < 0.01$).

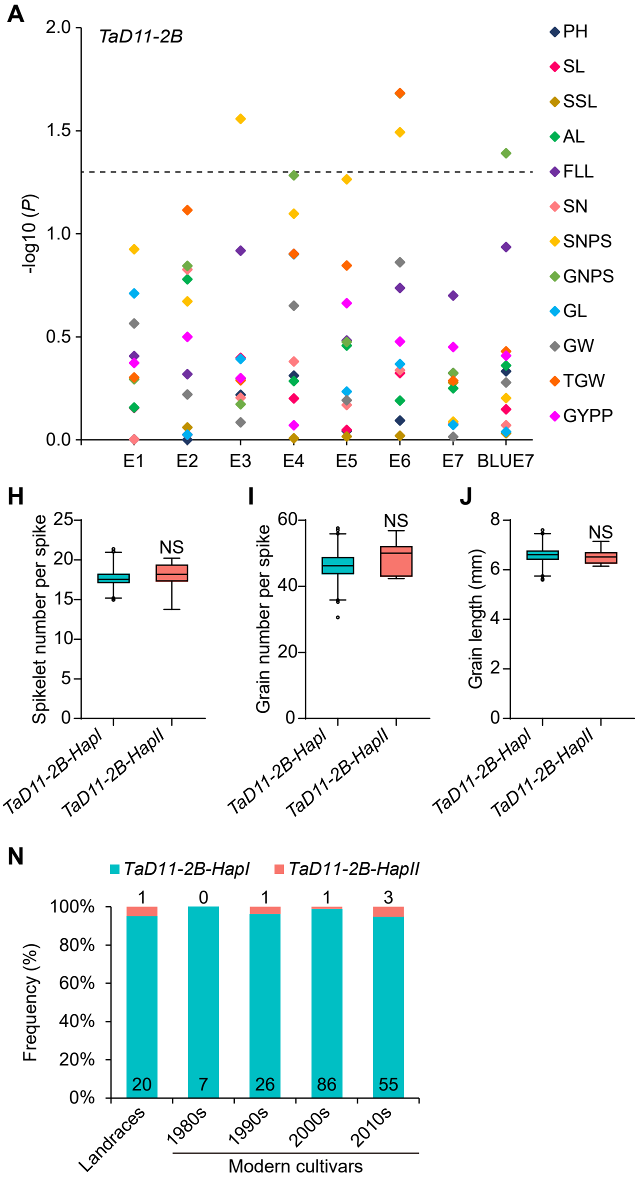


**Fig. S4.** Haplotype analysis of *TaD11-2B* associations with yield-related traits in wheat. (A) Genome-wide association analysis of *TaD11-2B* with 12 yield-related traits (PH, plant height; SL, spike length; SSL, spike stem length; AL, awn length; FLL, flag leaf length; SN, spike number; SNPS, spikelet number per spike; GNPS, grain number per spike; GL, grain length; GW, grain width; TGW, 1000-grain weight; and GYPP, grain yield per plant). E1–E7 represent association analysis in seven different environments. BLUE7 indicates association analysis based on the best linear unbiased estimate (BLUE) values calculated from seven environment data (E1–E7). Negative $\log_{10}$-transformed $P$ values are plotted. A black horizontal dotted line indicates the threshold value for significant associations ($P < 0.05$). (B–M) Comparison of 12 yield-related traits, including PH (B), SL (C), SSL (D), AL (E), FLL (F), SN (G), SNPS (H), GNPS (I), GL (J), GW (K), TGW (L), and GYPP (M) between *TaD11-2B* haplotypes based on BLUE values. Two-tailed Student's $t$ tests were performed between *TaD11-2B* haplotypes (NS = significant; $^{**}P < 0.01$). (N) Changes in the frequency of *TaD11-2B* haplotypes across the history of wheat breeding in China.

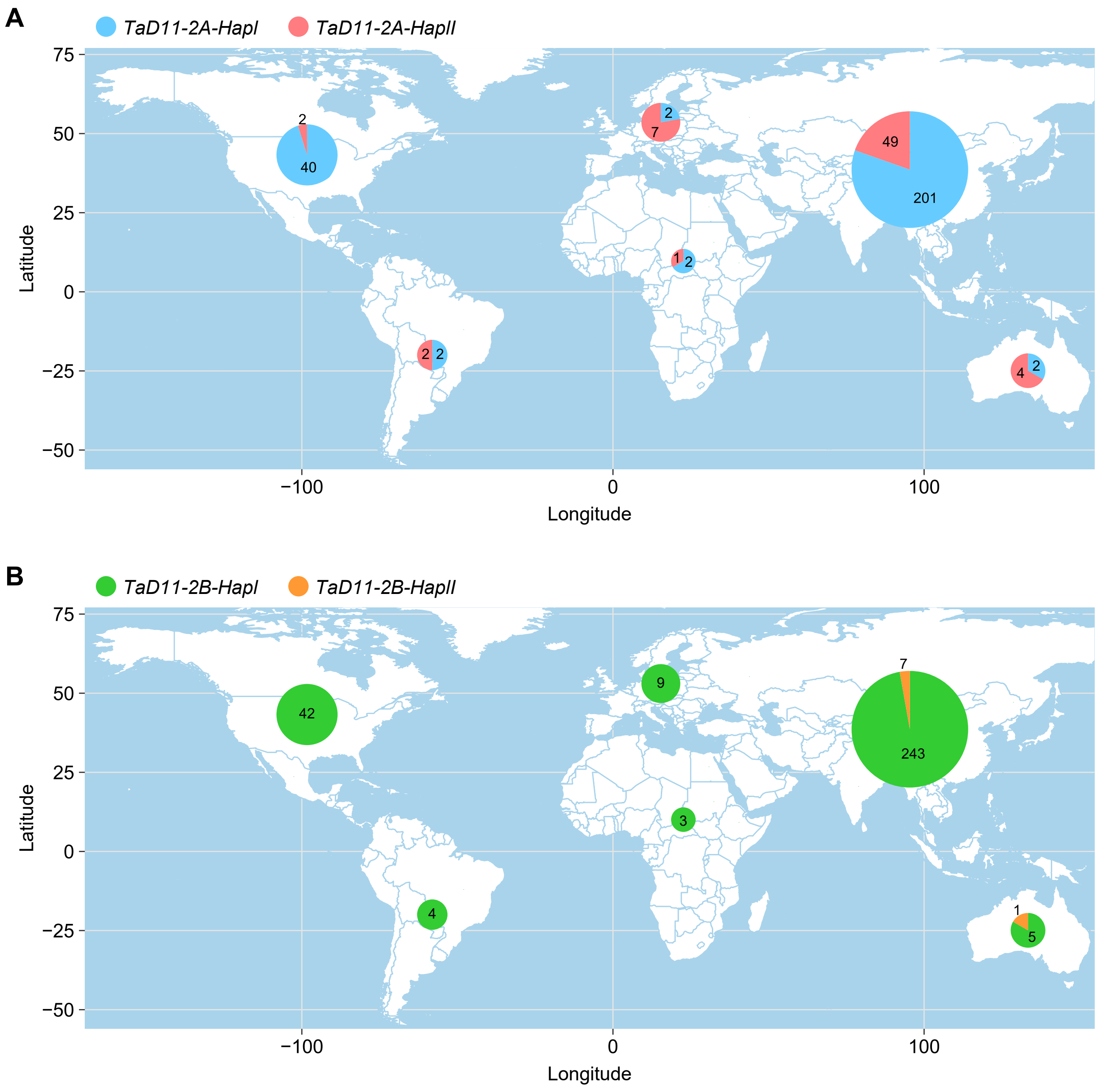


**Fig. S5.** Global distribution of *TaD11* haplotypes. (A) Distribution of *TaD11-2A* haplotypes across six continents. (B) Distribution of *TaD11-2B* haplotypes across six continents. There are 314 wheat accessions in this population, including 250 in Asia, three in Africa, nine in Europe, 42 in North America, four in South America, and six in Oceania.
